# *Timp2* loss-of-function mutation and TIMP2 treatment in murine model of NSCLC: modulation of immunosuppression and oncogenic signaling

**DOI:** 10.1101/2023.12.29.573636

**Authors:** David Peeney, Sarvesh Kumar, Tej Pratap Singh, Yueqin Liu, Sandra M. Jensen, Ananda Chowdhury, Sasha Coates-Park, Joshua Rich, Sadeechya Gurung, Yu Fan, Daoud Meerzaman, William G. Stetler-Stevenson

## Abstract

Mounting evidence suggests that the tissue inhibitor of metalloproteinases-2 (TIMP2) can reduce tumor burden and metastasis. However, the demonstration of such anti-tumor activity and associated mechanisms using *in vivo* tumor models is lacking. The effects of a *Timp2* functional mutation and administration of recombinant TIMP2 were examined in both orthotopic and heterotopic murine models of lung cancer using C57Bl/6 syngeneic Lewis Lung 2-luciferase 2 cells (LL2-luc2) cells. Mice harboring a functional mutation of TIMP2 (mT2) display markedly increased primary lung tumor growth, increased mortality, enriched vasculature, and enhanced infiltration of pro-tumorigenic, immunosuppressive myeloid cells. Treatment with recombinant TIMP2 reduced primary tumor growth in both mutant and wild-type (wt) mice. Comparison of transcriptional profiles of lung tissues from tumor-free, wt versus mT2 mice reveals only minor changes. However, lung tumor-bearing mice of both genotypes demonstrate significant genotype-dependent changes in gene expression following treatment with TIMP. In tumor-bearing wt mice, TIMP2 treatment reduced the expression of upstream oncogenic mediators, whereas treatment of mT2 mice resulted in an immunomodulatory phenotype. A heterotopic subcutaneous model generating metastatic pulmonary tumors demonstrated that daily administration of recombinant TIMP2 significantly downregulates the expression of heat shock proteins, suggesting a reduction of cell-stress responses. In summary, we describe how TIMP2 exerts novel, anti-tumor effects in a murine model of lung cancer and that rTIMP2 treatment supports a normalizing effect on the tumor microenvironment. Our findings show that TIMP2 treatment demonstrates significant potential as an adjuvant in the treatment of NSCLC.

**One Sentence Summary:** TIMP2 treatment compensates for TIMP2 mutation to reveal immunomodulatory and oncogene regulation that inhibits tumor growth and metastatic niche formation.

## INTRODUCTION

It is estimated that lung cancer will contribute to 12% and 21% of U.S. cancer cases and deaths in 2023, respectively (Surveillance, Epidemiology, and End Results (SEER) Program; (www.seer.cancer.gov)). Furthermore, the prevalence of lung cancer in low-income countries continues to rise, likely due to delayed and/or ineffective public health initiatives regarding the dangers of tobacco smoking, air pollution, and climate change (*1*). Lung carcinogenesis is accompanied by extensive changes in the composition and architecture of the pulmonary extracellular matrix (ECM), and the formation of an immunosuppressive tumor microenvironment (TME) (*2–6*). These changes can have a profound effect on disease progression, response to therapy, and incidence of relapse (*4–6*). Changes in the ECM disrupt contextual cues, both structural and compositional, that mediate normal homeostatic epithelial-mesenchymal interactions. For example, targeted expression of matrix metalloproteinase 3 (MMP3) in murine mammary epithelium results in an altered stromal environment that in turn promotes phenotypic conversion and malignant transformation of these epithelial cells (*7*). These epithelial-mesenchymal interactions have been shown to coordinate tissue organization and cellular identity in both developmental and pathologic conditions (*8, 9*). New strategies for lung cancer therapy are directed toward the TME and are the foundation for therapies to enhance host immune responses targeting malignant tumor cells (*10*). Developing strategies include the normalization of the microenvironment to support anti-tumor immune cell functions, reduce extracellular matrix turnover, and restore homeostatic mechanisms that prevent tumor progression (*11, 12*).

Tissue inhibitors of metalloproteinases (TIMPs) are endogenous inhibitors of metalloproteinase (MP) activity, with TIMP2 displaying the broadest expression profile of the family (*13, 14*). In addition to their functions as inhibitors of MP activity, TIMPs demonstrate a growing list of MP-independent functions that include the regulation of growth factor signaling, cytokine-like signaling capabilities, and direct modulation of cell behavior (*14–16*). We have previously reported that TIMP2 expression is a positive prognostic indicator in human breast and lung carcinomas (*17*). TIMP2 displays anti-tumor properties in murine xenograft as well as allograft models of lung adenocarcinoma and triple-negative breast cancer, respectively (*18–20*). These include suppression of tumor growth, tumor vessel density, and reduced vascular permeability, consistent with “vascular normalization” (*20*). In the present study, we developed a congenic C57BL/6j mouse line that harbors a large deletion within the *Timp2* gene, removing exons 2 and 3. This deletion results in diminished TIMP2 expression, significant reduction in MP inhibitory activity, and loss of the previously characterized α3β integrin-binding domain (*11, 14*). We characterized this new murine line and developed both orthotopic and heterotopic, syngeneic models of Lewis Lung Carcinoma to determine the effect of TIMP2 on the TME during lung tumor progression and the metastatic niche formation. We also investigated the effect of daily, systemic recombinant human TIMP2 (rTIMP2) delivery in tumor-bearing mice in a series of studies that highlight TIMP2 as a tumor-suppressor that functions within the tumor microenvironment of the diseased lung. Furthermore, we present evidence that TIMP2 acts as a “normalizing” factor within neoplastic tissues and represents a promising neo-adjuvant treatment strategy for solid tumors.

## RESULTS

### Generation of mutant *Timp2* (mT2) mice

A new congenic strain of mutant *Timp2* (mT2) mice containing a targeted mutation of the *Timp-2* locus (deletion of exons 2 and 3) on a clean C57/BL6j genetic background was generated by >10 generations of backcrossing of the original transgenic strain described by Catarina et al. (*21*), Figure 1A. These mutant mice lacking exons 2/3 were chosen specifically because this deletion results in the loss of the putative α3β integrin-binding domain of TIMP2 shown to mediate its MMP-independent anti-tumor effects, reviewed in (*11*). This targeted deletion disrupts Metzincin protease inhibition and loss of the α3β binding domain proposed to mediate many of the MP-independent effects of rTIMP2 *in vitro* and *in vivo* (*11, 22*). Deletion of exons 2 and 3 of the *Timp2* gene was confirmed by RNA sequencing and genomic PCR analysis, Figures 1B-D. The presence of both the wild type (wt, 283 bp) and mT2 bands (250 bp) in the same sample indicate that the mouse is heterozygous while single bands indicate either a homozygous wt or mT2, Figure 1D. Whole murine lungs from wt and mT2 mice were analyzed by immunoblotting for TIMP2 levels, indicating that mT2 protein levels are diminished compared to wt mice expressing full-length TIMP2 (Fig. 1E). Isolation of murine lung fibroblasts from wt and mT2 mice indicates that mT2 is poorly secreted and is principally found within the cell lysates, Figure 1F(i). Reverse zymography of the same samples reveals that mT2 has a diminished MMP inhibitory capability, as described previously (*21*), Figure 1F(ii). Characterization of the TIMP and gelatinase (MMP2/9) expression in the Lewis Lung Carcinoma cell line, LL/2-Luc2, grown as 3D spheroids reveals an appreciable expression of MMP2, TIMP2, and TIMP3, Figure 1G i and ii.

**Figure 1.**
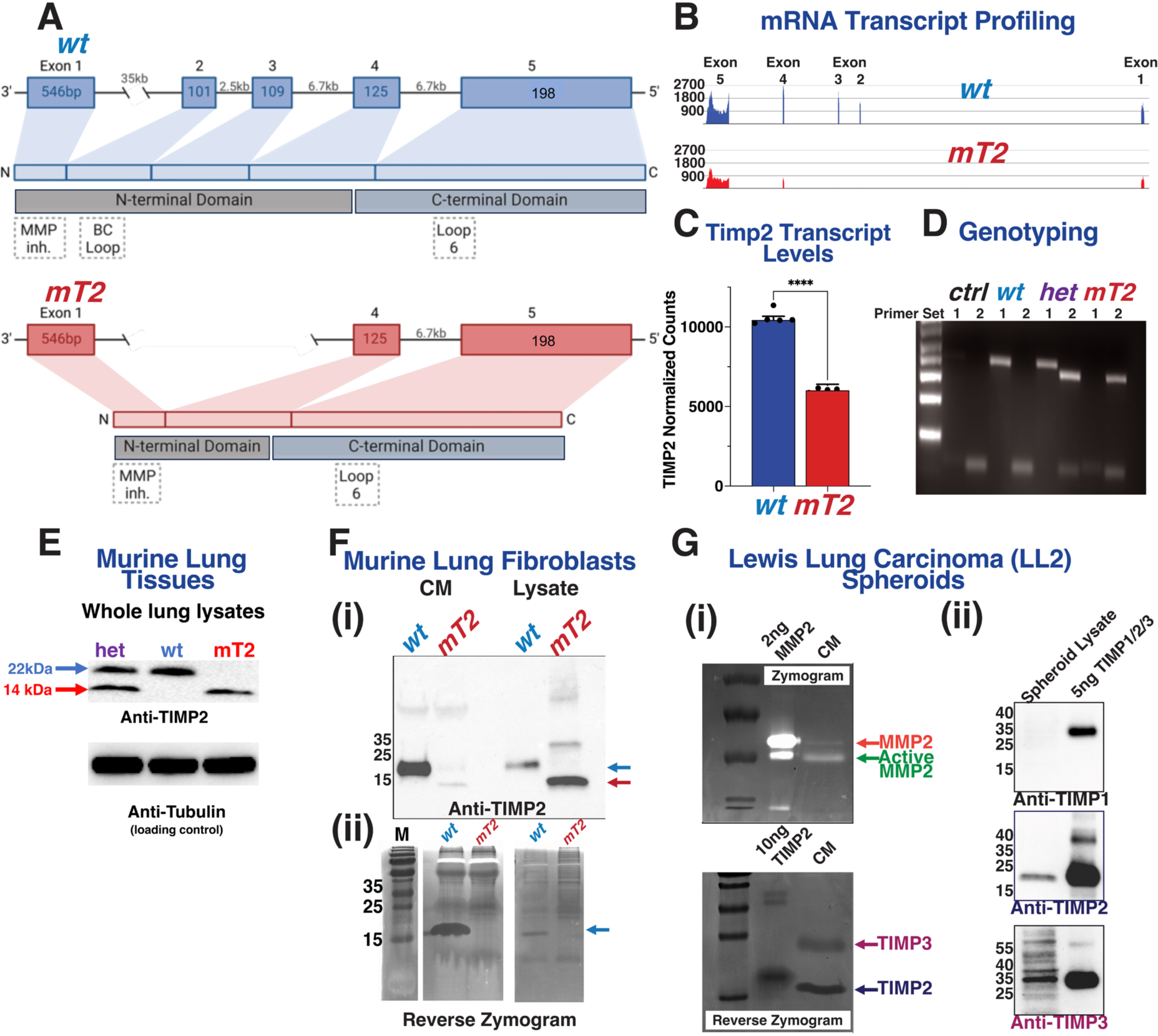
Characterization of the Lewis Lung Carcinoma Model. (A) Schematic describing the wild-type (wt) and mutant Timp2 (mT2) locus, and its translation into protein. (B) RNA sequencing confirms that the produced mutant Timp2 is lacking exons 2 and 3 and that this corresponds to a significant reduction in Timp2 transcript levels (C). (D) Example DNA genotyping using primers specific for wt (primer set 1) or mT2 (primer set 2). (E) Assessment of Timp2 protein levels in wt and mT2 lungs via immunoblotting. (F) Characterization of murine lung fibroblast Timp2 levels and activity through (i) immunoblotting and (ii) reverse zymography. (G) Characterization of Timp/gelatinase expression and activity of Lewis Lung Carcinoma (LL/2) cell spheroid conditioned media (CM) and lysates by (i) zymography, reverse zymography, and (ii) immunoblotting.

### Functional mutation of the *Timp2* gene increases tumor burden and mortality

To assess whether a loss of function mutation of the *Timp2* gene alters primary tumor growth, we employed an orthotropic murine model of Lewis lung carcinoma using the LL/2-Luc2 cell line. Using the experimental design outlined in Figure 2A, we compared overall survival, tumor growth, angiogenesis, and inflammatory responses in mT2 mice compared to wt mice. Kaplan-Meier analysis showed significantly higher mortality in mT2 mice relative to wt controls over 40 days, Figure 2B. In separate experiments, in vivo imaging measuring bioluminescence intensity (BLI) to quantify LL/2-Luc2 tumor growth showed a significant ∼10-fold increase in mT2 mice as compared to wt controls over a 28-day duration, Figure 2C. In an identical experiment using only female mice, we observed a ∼4-fold increase in tumor growth of mT2 compared to wt mice (Figure S1A & B). We also assessed the number of tumors by histology in the lungs of all mice and found that the number of tumor nodules was significantly higher (> 2-fold) in mT2 mice relative to wt, Figures 2D-E. In addition, the determination of the lung/body weight ratios, as a direct measure of pulmonary tumor burden, also demonstrates a > 2.5-fold increase in mT2 compared with wt mice. Determination of micro-vessel density (MVD) was performed in tumor tissues by immunohistochemical staining for CD31. A significant increase in tumoral MVD was observed in mT2 mice, Figures 2G i & ii), consistent with previous reports of TIMP2’s anti-angiogenic activity (*22*). Initial immunohistochemical staining of lung tumors from wt and mT2 mice demonstrates enhanced infiltration of Gr1^+^, CD45^+^, and CD11b+ myeloid cells in mT2 tumor-bearing mice, Figure 2H. Furthermore, immunofluorescence staining for the myeloid marker Gr1 showed enhanced positive staining in primary tumors, as well as the spleens from mT2 mice compared to wt mice, Figure 2H. This suggests that the functional mutation of *Timp2* results in enhanced production of myeloid-derived cells in organs of hematopoiesis (spleen) and an increase in their recruitment to tumor sites in wt mice. This led us to quantify the levels of myeloid cell subtypes in the lungs of both wt and mT2 mice, with or without orthotopic LL2-luc tumors, by flow cytometry, Figure 2I (gating strategies are shown in Figure S1C. These results demonstrate that there is a significant trend toward an increase in monocytes and neutrophils in non-tumor bearing mT2 mice compared with non-tumor bearing wt mice (*p* = 0.07 and *p* =0.14, respectively), which becomes statistically significant for the dendritic cell (DCs) population in non-tumor bearing mT2 mice compared with wt mice. These findings are consistent with tumor-independent activation of myeloid cell populations in non-tumor-bearing mT2 mice. Tumor-bearing wt mice did not demonstrate significant increases in any of these myeloid cell subtypes in comparison to non-tumor-bearing wt mice. However, tumor-bearing mT2 mice demonstrate increased levels of all three myeloid cell subtypes in comparison to tumor-bearing wt mice, and there is also a statistically significant increase in the neutrophil population of mT2 tumor-bearing mice compared with non-tumor-bearing mT2 mice. These findings are consistent with an altered myeloid cell response that is observed in both non-tumor- and tumor-bearing mT2 mice and may contribute to the enhanced tumor growth observed in mT2 mice. This effect would be mediated through the well-described effects of myeloid-derived cells in suppressing anti-tumor responses *in vivo*, including potent suppressive activity against effector lymphoid cells (*23, 24*). We also identified distinct immune cell foci in histologic sections of pulmonary tumors of both wt and mT2 mice, Figure 2 Ji. Quantification of the relative surface areas of these immune cell foci revealed significantly larger regions within mT2 tumors compared to wt controls, while differences in the number of inflammatory foci present were not statistically significant, Fig. 2Ji-iii. The histologic characteristics of these immune cell infiltrates showed that they were composed of small round cells with little cytoplasm and regular round nuclei consistent with lymphocyte morphology. Immunohistochemical staining revealed that these immune cell foci were CD3^-^, but positive for NKp46 staining, Figure 2Jiv. This suggests that these immune cell foci may represent aggregates of natural killer (NK) cells, that are not disseminated throughout the tumor. This finding suggests that the enhanced myeloid cell infiltrates observed in mT2 mice may result in the suppression of innate immunity as reported in many cancers (*23, 24*).

**Figure 2.**
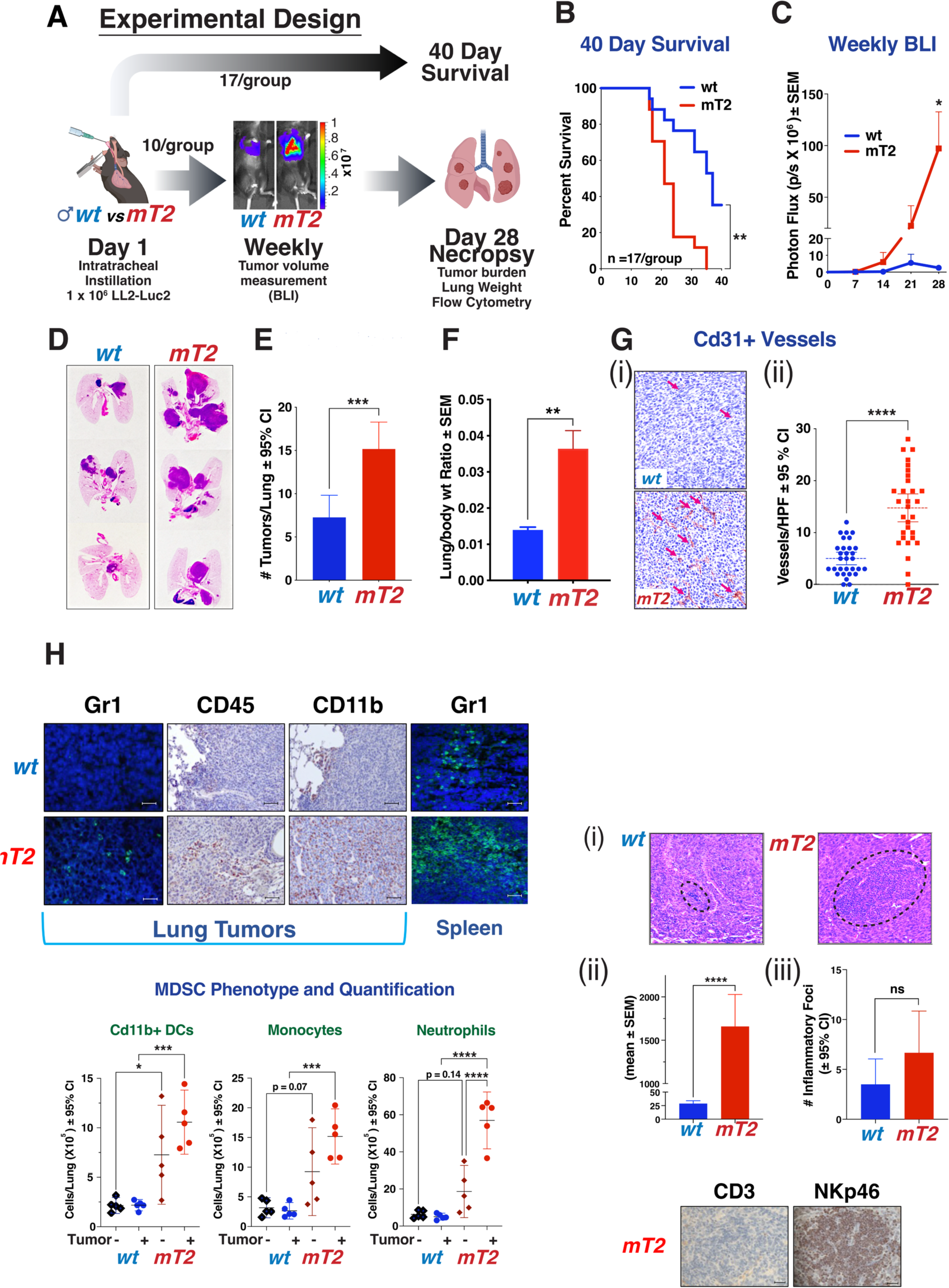
Mutant Timp2 Lewis Lung Carcinomas are phenotypically distinct from wt tumors. (A) The experimental protocol used for this series of experiments. (B) 40-day survival results of wt vs mT2 mice harboring orthotopic LL/2 tumors (*n* =17 mice/group). (C) BLI measurements of orthotopic LL/2 tumors (*n* = 10 mice/group). (D) Gross histological images of example lungs depicting significantly advanced tumors in mT2 mice. I Tumor counts and (F) tumor area of the H&E-stained lungs. (G) (i) Example immunohistochemistry and (ii) quantification illustrating the levels of Cd31+ vasculature within the primary tumors. (H) Immunostaining across wt and mT2 models for Gr1, CD45, CD11b. (I) Quantification of myeloid sub-types in whole lungs using flow cytometry. (J) (i) Example inflammatory foci, (ii) area, and (iii) quantification of which, within primary wt and mT2 tumors. (iv) Immunostaining of mT2 tumor foci for CD3 and NKp46.

### rTIMP2 treatment suppresses lung tumor growth

The loss of exons 2 & 3 in *Timp*2 causes significant decreases in the expression and protease inhibitory function of TIMP2, which in turn accelerates the tumor progression in mT2 mice. Next, we studied the effect of the administration of recombinant human TIMP2-6x-His (rTIMP2) in both mT2 and wt tumor-bearing mice. The TIMP2 protein sequence is highly conserved in mammals, with mature human and murine TIMP2 proteins sharing 98% sequence identity. To determine whether rTIMP2 could rescue the mT2 phenotype and suppress tumor growth, we performed a pilot experiment with a dose range of daily intraperitoneal injections (25, 50, 100, 200ug/kg/day) to characterize the dose-dependent response (Figure S1D and 1E. Although the tumor volumes as determined by BLI measurements in wt tumor-bearing mice were relatively low compared to the signals in mT2 tumor-bearing mice, a small but measurable reduction in tumor volumes was observed following rTIMP2 treatment, Figure S1E. In contrast, mT2 tumor-bearing mice demonstrated a significant reduction in tumor volumes at doses greater than 50 µg/kg/day in the same experiment. In addition, dosages of 100-200ug/kg/day rTIMP2 significantly improved 30-day survival in wt mice in a follow-up study, Figure S1F.

Based on these initial studies we examined the effects of systemic administration of TIMP2 on primary, orthotopic tumor growth in both wt and mT2 mice, as well as the transcriptomic changes that occur in whole mouse lungs harboring orthotopic Lewis Lung tumors in both wt and mT2 mice, with and without TIMP2 treatment. The experimental design for these experiments is outlined in Figures 3A and B. For both mT2 and wt, we compared transcriptomic analysis of non-tumor-bearing, tumor-bearing vehicle control (HBSS) treated, and rTIMP2-treated tumor-bearing mice. Tumors were initiated via intratracheal instillation as described previously (orthotopic, syngeneic tumor model) on day 1, with each group consisting of 5-7 mice (both male and female). Consistent with previous experiments, untreated mT2 mice exhibit accelerated tumor growth as demonstrated by BLI measurement of tumor volume and gross lung weight measurement of pulmonary tumor burden (Figure 3C). In this repeat experiment, the rTIMP2 treatment groups of both wt and mT2 mice (rTIMP2 at 200 µg/kg/day) demonstrate significant reductions in intrapulmonary tumor growth consistent with the observed effects in our initial dose-ranging experiments shown in Figure S1E. At the termination of the tumor growth experiment, the lungs were curated from all five mice in each group and prepared for RNAseq analysis of whole lungs (five mice per group for tumor-free, tumor-bearing, and rTIMP2-treated tumor-bearing mice).

**Figure 3.**
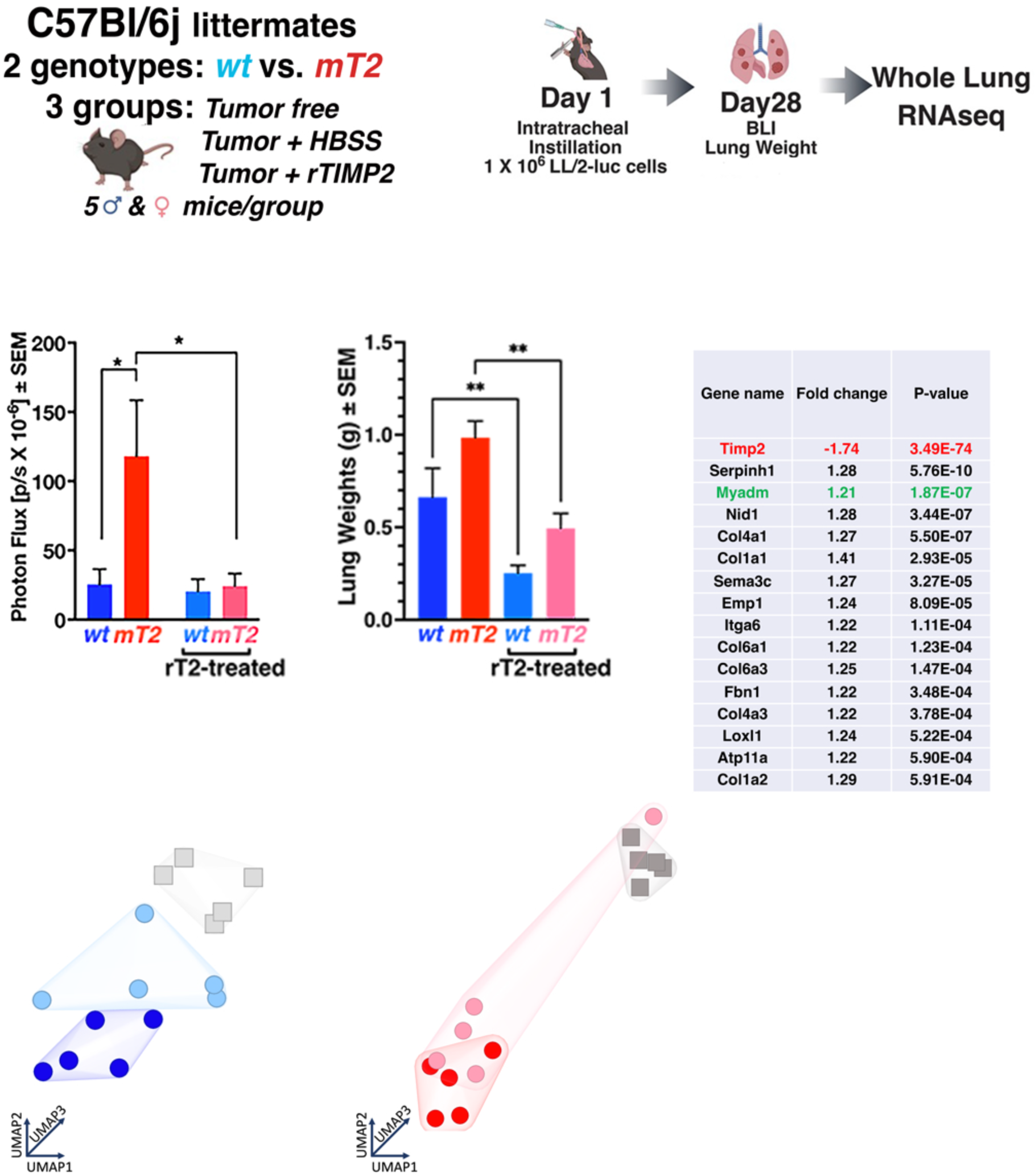
rTIMP2 inhibits the growth of orthotopic Lewis Lung Carcinomas. (A&B) Experimental design for the subsequent studies. I End-point tumor bioluminescence and lung weights. (D) List of 16 significant gene changes in tumor-free wt vs mT2 lungs. UMAP clustering of RNA sequencing data from I wt and (F) mT2 mouse lungs.

Considering the significantly enhanced tumor growth in mT2 compared with that observed in wt littermates, we anticipated profound alterations in the lung transcriptomes of non-tumor-bearing mT2 mice compared with wt littermates. Unexpectedly, using non-restrictive parameters (false discovery rate (FDR) 5 × 10^-2^, fold change cutoff (FC) > 1.2), we observed modest but significant changes in the expression of only 16 genes in comparing the transcriptome profiles of non-tumor-bearing lungs, Figure 3D. However, many of these changes affected widely expressed genes in the extracellular compartment, such as Col1a1, Col4a1, and fibronectin, and confirmed the reduced expression of *Timp2*, as demonstrated in our initial characterization of these mT2 mice. In addition, modest changes in several genes associated with enhanced tumor growth and progression, such as collagen VI (*Col6a1* and *Col6a3*), ATPase Phospholipid Transport-11a (*Atp11a*) and epithelial membrane protein 1 (*Emp1*), as well as a modest increase in expression of the Myeloid-associated Differentiation marker (*Myadm*), a reported negative prognostic biomarker in NSCLC (*25–29*). However, this analysis did not reveal significant changes in the expression of other *Timp* or *Mmp* genes but did reveal a slight increase in the expression of the serine protease inhibitor *Serpinh1*.

UMAP analysis of wt tumor-bearing mice demonstrates distinct segregation of the rTIMP2 treatment group from both the tumor-free and vehicle control (HBSS) groups, indicating that rTIMP2 treatment induces a distinct transcriptomic profile in tumor-bearing lungs, Figure 3E. This rTIMP2-treated group of wt mice is juxtaposed directly between the vehicle control tumor-bearing mice (HBSS) and tumor-free wt mice (Figure 3E). These data suggest that the gene expression profiles of rTIMP2-treated tumors in wt mice are distinct from those of tumor-bearing vehicle control treated (HBSS) mice. A similar analysis of tumor-bearing mt2 mice failed to demonstrate distinct clustering or separation of the rTIMP2-treated mice from the vehicle control group (Figure 3F), except for a single outlier. This finding suggests that, despite the clear effect of rTIMP2 on primary pulmonary tumor growth in this cohort (Figure 3C), rTIMP2 treatment had a markedly different impact on the transcriptome profile of these tumor-bearing lungs. As a result, further analysis of RNA sequencing data from these experiments was separated by genotype to focus specifically on the effects of exogenous rTIMP2 in modulating tumor growth separately in wt and mT2 mice.

In wt lungs, utilization of a stringent FDR threshold of ≤ 1 x 10^-4^ reveals a cluster of 221 genes that display significant alterations in expression across two combined parameters; (1) vehicle control versus tumor-free and (2) vehicle control versus rTIMP2-treated, summarized in the hierarchical clustering diagram in Figure 4A. Euclidean distance was employed to produce the cluster dendrogram, and this shows that 80 % of mice in the rTIMP2 treatment group show gene expression patterns that cluster with the non-tumor-bearing mice (Figure 4A). In mT2 lungs, equivalent analysis with a relaxed FDR (≤ 5 x 10^-2^) for the rTIMP2-treated versus vehicle control comparison highlights 236 gene changes. Cluster analysis (Euclidean) reveals 3 major clusters, with 80 % of the rTIMP2-treated tumor-bearing mT2 mice forming a unique cluster that is distinct from the tumor-free group but overlaps with 2 out of 5 vehicle control (HBSS) treated tumor-bearing lungs (Figure 4B). Relaxing the FDR (≤ 5 x 10^-2^) for the wt rTIMP2-treated versus vehicle control comparison increases the gene list to 2020, which reveals a profound “normalization” of tumor-associated gene changes in wt lungs when the fold changes across each treatment group are plotted (Figure 4A and C). In the mT2 tissue samples, there were 236 significant gene changes induced by TIMP2 treatment (FDR ≤ 5 x 10^-2^). To produce a fold change plot with comparable gene numbers to the corresponding analysis for wt lungs (Figure 4C), we utilized only the tumor-associated gene changes (vehicle control versus tumor-free, FDR ≤ 1 x 10^-4^) which equates to 1825 genes. These analyses unveil a distinct “normalizing” pattern across the transcriptome of rTIMP2-treated tumor-bearing lungs that is most prominent following rTIMP2 treatment in wt mice, Figures 4A and C. We refer to this effect as normalizing in the sense that the data in both formats demonstrate that the effect of rTIMP2 treatment results in transcriptome changes that are much more similar to those observed in wt tumor-free mice. In the experiments examining the effects of rTIMP2 treatment in mT2 mice, the analysis of fold changes for these 1825 tumor-associated genes reveals a less pronounced, but significant shift in the transcriptomic signature that is most pronounced in the genes that are downregulated in the tumor-bearing, HBSS (vehicle control) treated mice, Figures 4B and D. Whereas, in wt mice changes are observed in genes with both enhanced and reduced expression, Figure 4A and C.

**Figure 4.**
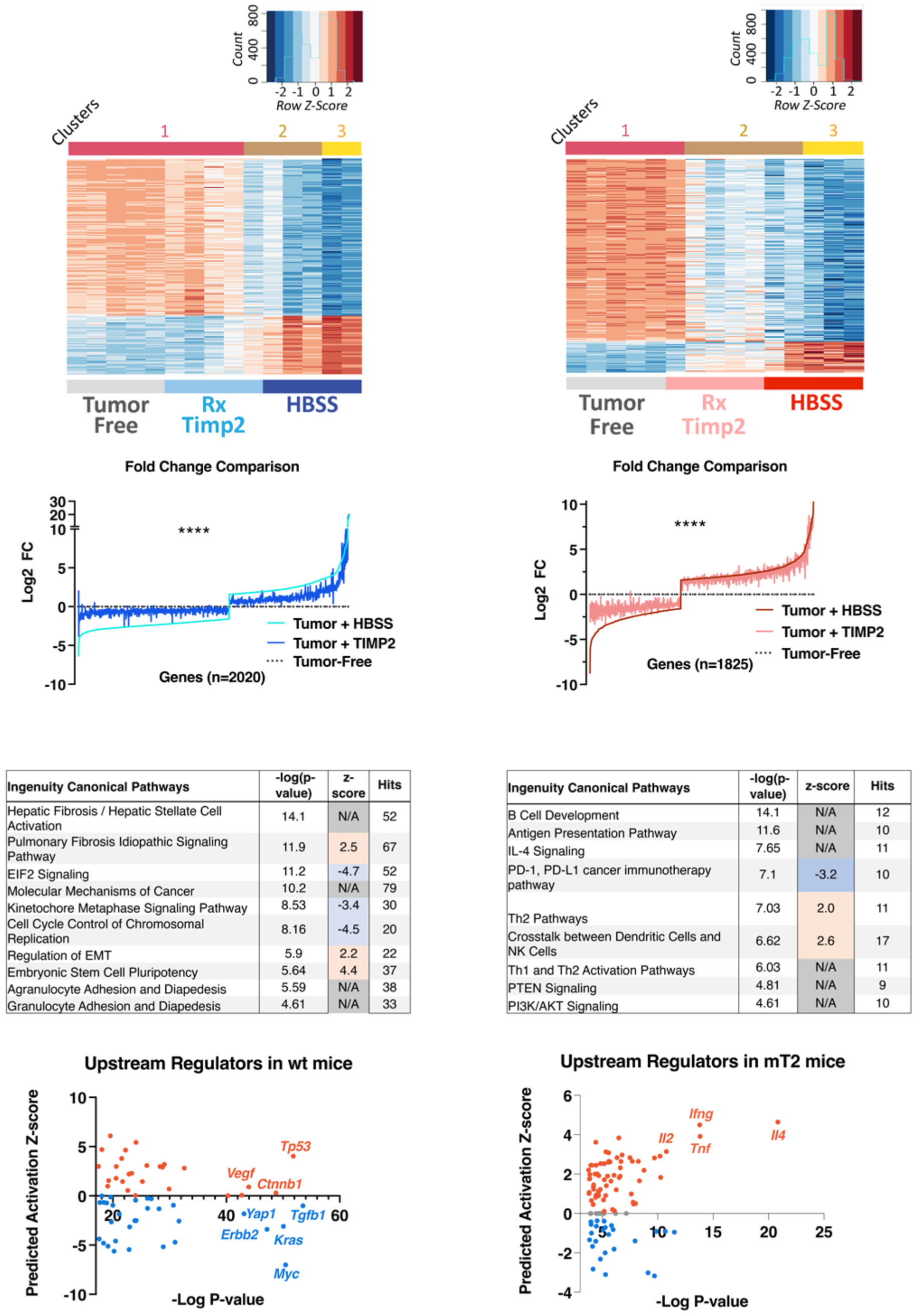
Recombinant TIMP2 treatment in tumor-bearing mice supports the normalization of tissue transcriptome. (A&B) Heatmap illustrating relative expression of differentially expressed genes across all treatments in (A) wt and (B) mT2 mice. (C&D) Comparison of the Log2 fold change (FC) across all treatments in (C) wt and (D) mT2 mice. (E&D) Top significantly altered molecular pathways identified by Ingenuity Pathway Analysis in rTIMP2-treated wt I and mT2 (F) mice, with the associated plotted upstream regulators for wt (G) and mT2 (H) data.

Gene set pathway analysis further highlights the differential responses following rTIMP2 treatment of these intrapulmonary tumors in wt compared with mT2 mice, Figure 4E and F, respectively, with the tabular data sets included in Table S1. In wt mice, the rTIMP2-treated tumor-bearing lungs demonstrate suppression of the cell stress response associated with the Eukaryotic Initiation Factor 2 (*EIF2*) signaling pathway, as well as downregulation of pathways associated with cell proliferation such as the Kinetochore Metaphase and Cell Cycle Control of Chromosomal Replication pathways. This analysis also predicts upregulation of the Pulmonary Fibrosis Idiopathic signaling pathway, however, no evidence of increased pulmonary fibrosis was observed during the histologic examination of these tumors. Prediction of upstream regulators controlling the observed gene changes suggests reduced activity of several oncogenic regulators such as *Myc*, *Kras*, *Tgfb1*, *Erbb2*, and *Yap1*, as well as increased activity of *Tp53* tumor suppressor gene (Figure 4G). These findings sharply contrast with the analysis of the 236 gene changes identified in response to rTIMP2 treatment in tumor-bearing mT2 mice, Figure 4F. The canonical pathways and upstream regulators identified by gene set enrichment using Ingenuity Pathway Analysis reveal a unique signature for rTIMP2-treated tumors in mT2 mice compared with vehicle control treatment (Figure 4F & 4H). Surprisingly, the effect of rTIMP2 in mT2 tumor-bearing mice is associated with the downregulation of the PD-1, PD-L1 immunoregulatory pathway. This pathway is the target of immune checkpoint inhibitory therapy that has significantly improved responses in cancer therapy. In addition, this analysis also suggests that rTIMP2 treatment results in the activation of Th2 cells and enhanced crosstalk between dendritic cells and NK cells, which are both associated with immune cell-mediated anti-tumor activity. The Upstream regulators identified in these analyses suggest enhancement of *Il4*, *Ifng, Tnf*, and *Il2* activity and down-regulation of *Il10*, Figure 4H. These changes are consistent with a complex pattern of immune regulation that promotes enhanced T-cell function combined with down-regulation of acute inflammatory responses. These findings show that the effect of rTIMP2 treatment is dependent on the genetic background of the tumor model. In mT2 tumor-bearing mice, the response to rTIMP2 treatment facilitates immune modulation, whereas in wt mice rTIMP2 treatment mediates suppression of cell stress and mitogenic pathways. These findings suggest that rTIMP2 treatment in mT2 mice may remove the checkpoint inhibition mediated by enhanced MDSC activation in this genotype. As a result, we posit that enhanced T-cell responses and innate immune cell anti-tumor activities may mediate the beneficial effects of rTIMP2 treatment in tumor-bearing mT2 mice. This proposed immunomodulatory role for TIMP2 in reversing an immunosuppressive, pro-tumorigenic environment is consistent with prior reports demonstrating TIMP2 alteration of myeloid responses and NK cell activity (*30, 31*).

Irrespective of this apparent lack of similarity in the overall patterns of gene expression in the analysis of lungs from rTIMP2-treated wt and mT2 mice, there is a distinct subset of 119 similar gene changes between the two data sets, Figure 5A, Table S2. Gene set analysis of the 119 similar genes between the wt and mT2 datasets reveals a possible central role for rTIMP2 treatment in modulating immune cell function which centers around changes in *Tnf*, *IL4*, and Vegf activity, as shown in Figure 5B. These data suggest that although more prominent in the mT2 data sets, the response following rTIMP2 in both mT2 and wt mice implicates immune modulation mediating an improvement in innate immune (T and NK cell) responses, and an enhanced anti-tumor immune response.

**Figure 5.**
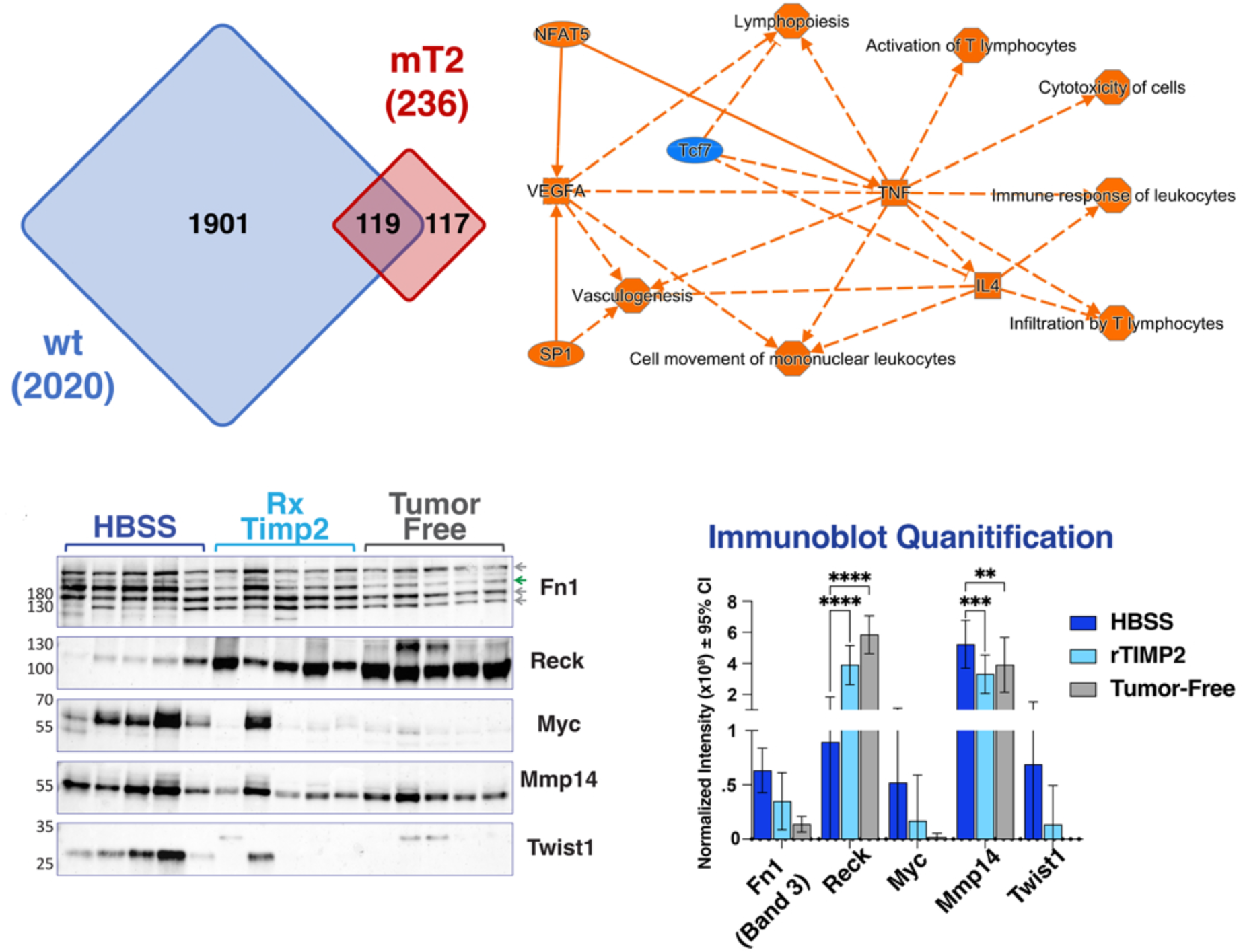
Comparison of recombinant TIMP2 treatment in wt and mT2 mice. (A) Venn diagram depicting the unique and shared transcriptome changes in rTIMP2-treated tumor-bearing wt and mT2 mice. (B) A graphical summary from Ingenuity Pathway Analysis of the 119 shared genes between the wt and mT2 models. (C&D) Immunoblotting, and quantification of which, of a selection of significant differentially expressed genes across all treatments.

We also examined the effects of rTIMP2 treatment on protein expression levels of select markers in whole lung tissue extracts of wt tumor-bearing (HBSS and rTIMP2-treated) as well as non-tumor bearing (tumor-free) mice (5 mice/group), Figure 5C and D. Quantification of these immunoblot results demonstrate a trend in reduced expression of several oncogenic markers, Fibronectin 1 (Band 3, FN1), Myc, MMP14 and the epithelial-to-mesenchymal transition (EMT) marker Twist. In contrast, this analysis also demonstrates increased protein expression of the reversion-enhancing, cysteine-rich protein with Kazal motifs (RECK). RECK is a membrane-anchored glycoprotein with multiple epidermal growth factor (EGF)-like repeats, as well as serine protease inhibitor-like domains (*32, 33*). RECK also has demonstrated MMP inhibitor activity, and its expression is associated with loss of the invasive, metastatic phenotype. Quantification of the immunoblot results demonstrates a statistically significant increase in RECK expression and reduction of MMP14 in rTIMP2 treated wt tumor-bearing mice compared with vehicle control (HBSS) treated wt tumor-bearing mice. The results show these changes are similar to basal levels observed in tumor-free mice. These findings are consistent with the rTIMP2-treatment-mediated reduction in oncogenic drivers observed in the RNAseq analysis comparing vehicle control treated wt tumor-bearing lungs with rTIMP2-treated tumor-bearing lungs of wt mice. These findings are also consistent with previous reports documenting the reduction of mesenchymal transcription factors, such as Twist, and enhanced RECK expression in human tumor cell lines following rTIMP2 treatment *in vitro* (*11, 18*).

### Treatment with rTIMP2 alters the metastatic niche formation in wt mice

In the present study, we utilized a heterotopic, syngeneic model of spontaneous metastasis (Lewis Lung cancer model) to examine the effect of rTIMP2 treatment on the metastatic tumor microenvironment in the lungs of wt C57Bl/6 mice bearing subcutaneous, LL2-luc2 dorsal flank tumors, Figure 6 and Figure S2 (*34*). In this preliminary experiment, we subcutaneously inoculated 1 × 10^6^ LL2-luc2 cells in the lateral dorsal flank region of 24 wt C57Bl/6 male mice. At subsequent, regular 7-day intervals (days 7, 14, 21, and 28) we randomly selected and euthanized 6 mice and evaluated pulmonary metastatic burden (total lung weights), as described in Figures S2A and B. This preliminary experiment revealed a significant increase in total lung weights between days 7 and 21 that reflects an increase in pulmonary metastasis. This correlation between total lung weight and bioluminescence/histologic tumor burden has been demonstrated in our previous experiments, Figures 2 and 3, and correlates with an increase in the levels of luc2 gene expression observed (see below, Figure 6Cvi). These findings are consistent with previous reports utilizing spontaneous metastasis models of Lewis Lung carcinoma in that the lungs are the predominant site of metastasis from these subcutaneous, dorsal flank tumors (*34, 35*).

**Figure 6.**
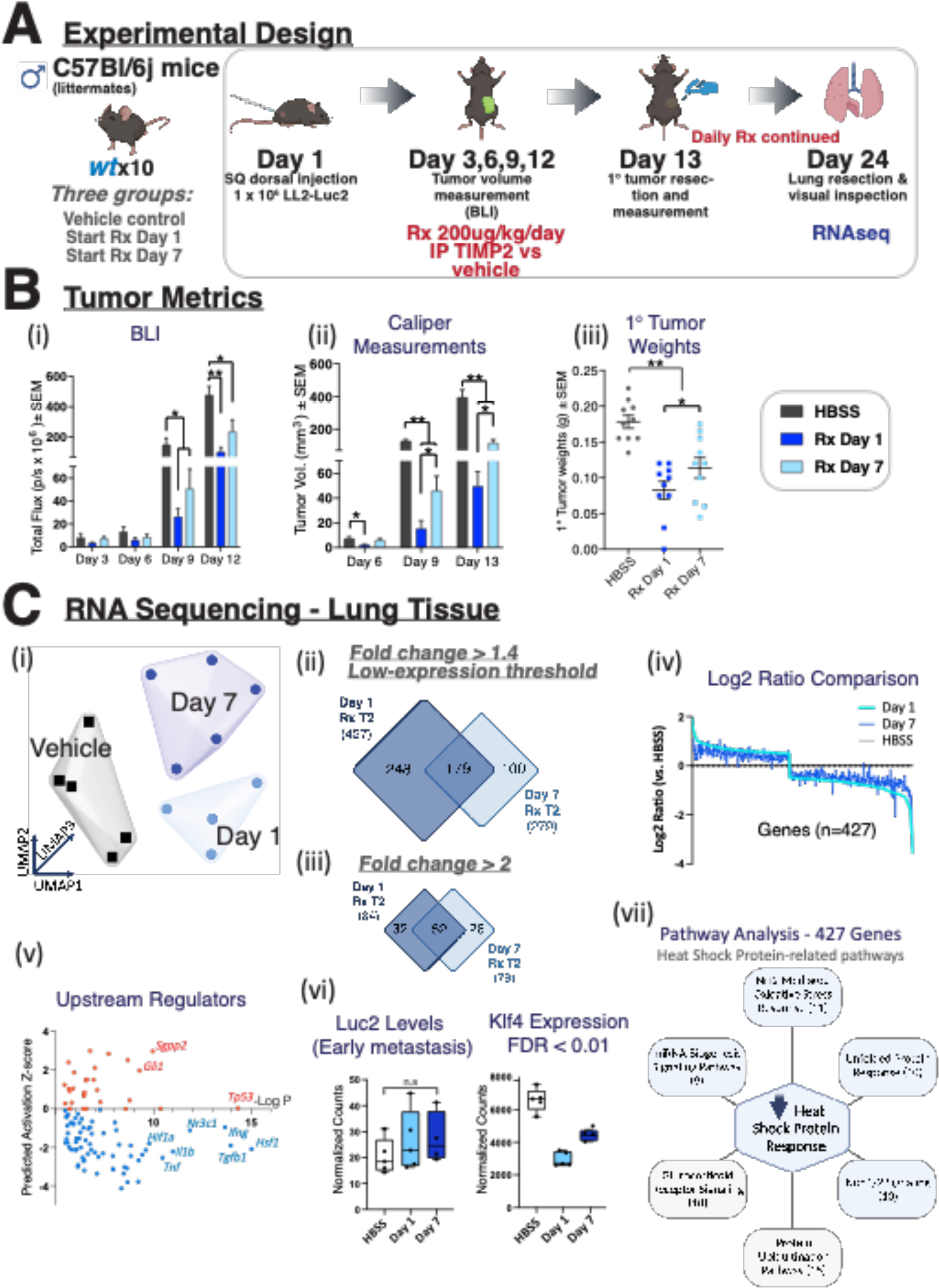
rTIMP2 treatment inhibits subcutaneous LL/2 tumor growth and produces a unique transcriptional profile in metastatic tissues. (A) Schematic describing the experimental design used in this study. (B) Tumor metrics describing (i) bioluminescence imaging, (ii) caliper measurements, and (iii) resected weights of the subcutaneous primary LL2-Luc2 tumors. (C) RNA sequencing of lungs reveals that rTIMP2 promotes a unique transcriptome, evidenced through (i) UMAP clustering and differential expression analysis. (ii + iii) Venn diagrams depicting the unique and shared gene changes between both treatment regimens, segregated by analysis parameters. (iv) Comparison of the log2 ratio (rTIMP2 vs HBSS) reveals a similar expression profile between the two rTIMP2 treatments. (v) Plot illustrating the top 100 upstream regulators identified through Ingenuity Pathway Analysis that likely display an alteration in activation status. (vi) Luc2 gene expression indicates that these lungs are early metastatic, yet rTIMP2 treated mice display reduced expression of the pre-metastatic niche marker Klf4. (vii) Pathway analysis highlights a clear alteration in the Heat Shock Protein Response, which is heavily involved in the pathways listed as significantly altered through Ingenuity Pathway Analysis. The number under each pathway identifies the number of gene hits in the designated pathway from the rTIMP2 treatment schedule that started on day 1.

In subsequent experiments, we examined the effect of rTIMP2 administration on the pulmonary transcriptomic profiles of wt mice following implantation of heterotopic, subcutaneous, dorsal flank tumors. As outlined in the Experimental Design (Figure 6A), treatments with rTIMP2 were initiated in wt groups either on the day of tumor initiation (Day 1) or seven days after tumor initiation (Day 7), with resection of primary subcutaneous tumors on day 13, to mimic neoadjuvant treatment strategies. In these experiments, rTIMP2 effectively reduced primary tumor growth by greater than 50% when measured through in vivo imaging and caliper measurements in both treatment groups (Day 1 and Day 7), Figure 6Bi-ii. Measurement of resected tumor weights following primary tumor resection (Day 13) shows that the rTIMP2 suppressive effect on tumor growth was slightly greater in the treatment group initiated on Day 1 compared to the start of treatment on Day 7, Figure 6 B. Comparison of primary subcutaneous tumor weights at the time of resection on day 13 shows that rTIMP2 treatment, starting on Day 1 or Day 7, resulted in 50 to 35 % reduction in tumor weight, respectively (Figure 6B iii). Following primary tumor removal on day 13, daily rTIMP2 treatment via IP injections was continued. The mice were euthanized, and lungs were collected for analysis on Day 24. The lungs displayed no grossly observable tumor formation upon visual inspection.

The lungs were subjected to bulk RNA sequencing analysis to assess for micro-metastasis and global gene expression. The results confirmed that the burden of pulmonary metastasis was very low in the control and rTIMP2-treated mice with no significant difference between the three groups of mice, see below. UMAP analysis of the normalized sequencing data reveals a unique gene expression profile of vehicle control treated wt tumor-bearing mice compared to lungs from rTIMP2-treated tumor-bearing wt mice with rTIMP2-treatment started on day 1 and day 7, Figure 6 Ci. Differential expression analysis with a low fold change cut-off (1.4) and minimum expression cut-off (200 normalized counts) revealed 427 and 279 significant gene changes for the rTIMP2-treated tumors starting day 1 and day 7, respectively, Figure 6Cii. Increasing the fold change cut-off to 2.0 and removing the minimum expression threshold reduces the number of identified gene changes, but also identifies additional, significantly altered, low-expressed genes, Figure 6Ciii, all of which are identified in Table S2. The transcriptome changes following rTIMP2 treatment in both treatment schedules (starting day 1 and day 7) include a consistent reduction in numerous heat shock proteins, including members of the Hspb, Hsp10, Hsp40, Hsp60, Hsp70, Hsp90, and Hsp110 families (See Table S2 for details). Comparison of the calculated log2 ratios for each rTIMP2 treatment schedule reveals a clear correlation between lung transcriptome profiles, Figure 6Civ. This would suggest that over time the delayed start of treatment could be as effective as the start of treatment on Day 1. The identification of potential upstream regulators of the gene changes observed following rTIMP2 treatment of wt mice with lung metastasis suggests significant upregulation of *Tp53* and suppression of *Tgfb*, analogous to changes observed in wt mice bearing primary lung tumors. However, we also observed a downregulation of *Hsf1* that was seen in prior experiments involving primary lung tumors, Figure 6v. Across all samples, we find similar levels of Luc2 gene expression, suggesting that the burden of micro-metastases in these experiments was similar between rTIMP2-treated and vehicle control-treated mice, Figure 6Cvi. Despite no clear difference in metastatic burden in the lungs of rTIMP2-treated and control-treated mice, we observed distinct changes in the overall pattern of gene expression that included a significant downregulation in Klf4 expression, a reported pre-metastatic niche marker, Figure 6Cvi, (*36*). Pathway analysis demonstrates downregulation in multiple genes that are involved in distinct biological pathways that coalesce around the suppression of heat shock responses, Figure 6Cvii. Upstream analysis of the 427 significant genes highlighted the downregulation of master regulators implicated in both the heat shock response (*Hsf1*) and inflammation (*Tgfb1*, *Tnf, Il1b*, *Igf1*), Figure 6Cv, highlighting that the essential differences between rTIMP2 treatment schedules are quantitative rather than qualitative and that the delayed initiation of treatment, representative of typical clinical interventions, may still provide therapeutic benefit.

## DISCUSSION

Often described solely as an inhibitor of metalloproteinases, there is a growing appreciation of the MP-independent functions of TIMP family proteins and their diverse roles in tissue biology. The MP-independent effects of rTIMP2 in suppressing the proliferative and invasive properties of a variety of cell types, which are mediated via the α3β integrin/cAMP/PKA/Shp1/p27-dependent pathways and resulting in enhanced nuclear p27 localization, enhanced Rap1 and p53 signaling have been reviewed extensively elsewhere (*11, 14*). Despite few reports to the contrary (*37–39*), growing evidence suggests that TIMP2 principally displays anti-tumor capabilities in a manner that is not limited to its role in MP inhibition (*19, 20, 30, 40–42*). We have previously shown that rTIMP2 can reduce tumor burden and metastasis in several *in vivo* tumor models (*19, 20, 31, 43*). The Lewis Lung Carcinoma tumor model is widely used and considered an accurate representation of human adenocarcinomas, providing the foundation of numerous discoveries in lung cancer progression (*34, 44, 45*).

In this study, we present evidence describing the anti-tumor and anti-metastatic capabilities of the matrisome protein TIMP2, using orthogonal approaches. We first developed and characterized a congenic C57Bl/6 murine line carrying loss-of-function mutation through deletion of TIMP2 exons 2 & 3, from the previously described mutant TIMP2 model described by Caterina and colleagues (*21*). In our initial set of experiments, we characterized the growth of orthotopically implanted Lewis lung carcinoma cell tumors in syngeneic wt and mT2 mice, summarized in Figure 7A. The results demonstrate a significant enhancement of tumor growth and diminished survival of mT2 tumor-bearing mice compared with wt littermates inoculated with the same initial tumor cell burden. In addition, the lung tumors present in mT2 mice were more vascularized and demonstrated a more robust infiltration of myeloid-derived suppressor cells (MDSCs) as evidenced by enhanced immunostaining for myeloid markers Gr1, CD45, and CD11b, as well as flow cytometric quantification of specific myeloid cell subpopulations. A detailed analysis of the increase in myeloid subpopulations, showing an increase in both tumor-free and tumor-bearing mT2 mice, has not previously been fully characterized.

**Figure 7.**
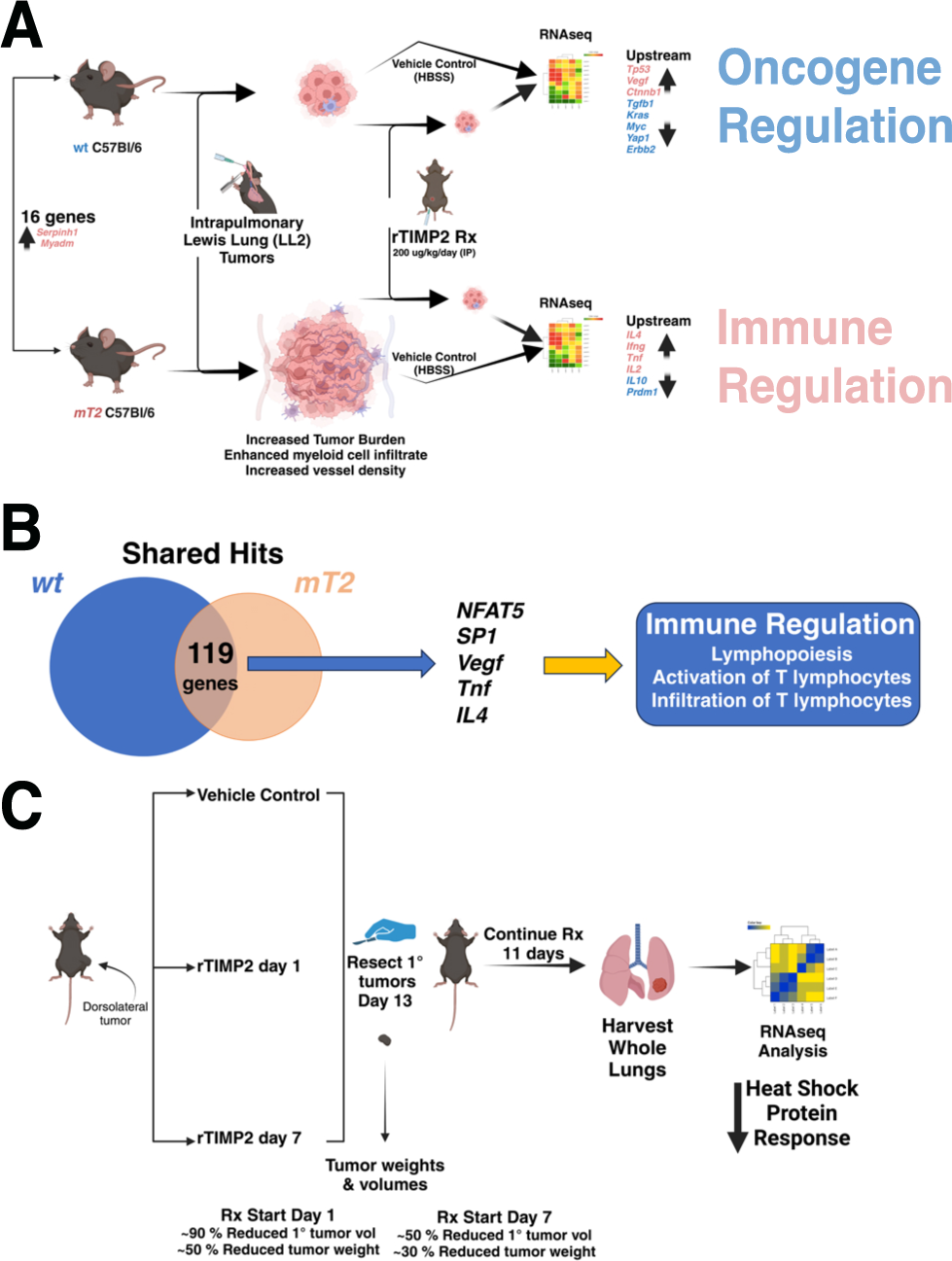
Summary and Conclusions. (A) Summary of primary lung tumor experiments with both wt and mT2 mice. Comparison of transcriptional changes between genotypes reveals changes in expression of 16 genes, including modest upregulation of *Myadm*. The inoculation of primary lung tumors demonstrates enhanced tumor growth in mT2 mice, but daily treatment with rTIMP2 reduced primary lung tumor growth in both wt and mT2 mice. RNA seq analysis demonstrates distinct patterns of gene expression in wt and mT2 mice. Tumor-bearing, rTIMP2 treated wt mice gene changes are consistent with oncogene regulation, whereas in mT2 the response suggests an immunoregulatory phenotype. (B) A summary of shared hits between the wt and mT2 phenotypes described in panel A, reveals an underlying commonality of an immunoregulatory phenotype of enhanced lymphopoiesis along with enhanced activation and infiltration of T lymphocytes, not previously associated with TIMP2 treatment. (C) Summary of study examining the effects of rTIMP2 treatment on the metastatic niche in wt mice. RNAseq analysis of whole lungs from rTIMP2-treated and control-treated lungs shows downregulation of the heat shock protein response.

Furthermore, our initial experiments identified immune cell foci that while present in both mT2 and wt mice were larger in lung tumors of mT2 mice. Immunohistochemical staining of these immune cell foci suggests that they were principally composed of NK cells. These readily identifiable immune cell clusters are consistent with prior reports describing the MDSC suppression of innate immune cell function, including CD8^+^ and NK cells, resulting in enhanced tumor growth and metastasis (*46, 47*). These findings strongly suggest that further investigations of the effects of the functional mutation of TIMP2 on innate immune cell function in lung cancer are warranted but beyond the scope of the present study.

In our second set of experiments, we again examined the growth of primary lung tumors in wt and mT2 mice but included a comparison of the treatment effects on tumor-bearing mice with either rTIMP2 or vehicle control (HBSS) for 28 days, Figure 7A. In addition, whole lung tissues from tumor-free, rTIMP2-treated, and vehicle-control treated mice from both the wt and mT2 littermate cohorts were processed for RNAseq analysis. First and foremost, rTIMP2 treatment reduced tumor burden in both wt and mT2 mice, consistent with previous reports that TIMP2 demonstrates significant anti-tumor activity (*18–20, 46, 48*). Surprisingly, a comparison of RNAseq data from wt and mT2 tumor-free mice revealed modest (less than two-fold), but significant changes in the expression of 16 genes. These changes include the anticipated decrease in *Timp2* expression, but slight increases in genes associated with enhanced myeloid cell activation (*Myadm*), as well as enhanced tumor growth as described above. Based on these findings, we conducted separate analyses of rTIMP-treated and non-treated lung tumor-bearing mT2 and wt mice RNAseq data.

Transcriptome profiling of tumor-free and tumor-bearing mice with and without rTIMP2 treatment initially reveals distinct profiles for wt and mT2 mice, Figure 7A. rTIMP2-treated tumor-bearing wt mice exhibit gene expression changes that focus on the downregulation of oncogenic pathways such as EIF2 signaling and cell cycle control of chromosomal replication, a result of reduced activity of upstream oncogene drivers such as *Myc*, *Kras*, and *Tgfb*, and enhanced activity of the tumor suppressors *p53* and *Ctnnb1* (Beta-catenin). In contrast, the transcriptional changes in rTIMP2-treated tumor-bearing mT2 mice suggest that there is suppression of the inflammatory response and a corresponding enhancement of the innate immune response as evidenced by suppression of the PD-1, PD-L1 checkpoint pathways, and enhancement of crosstalk between dendritic cells and NK cells. There is also an enhancement of the Th2 pathway, consistent with the identification of an increase in *Il4* activity observed in an analysis of upstream regulators. Between the wt and mT2 models, there were 119 similar gene changes with an FDR of 5 x 10^-2^. Analysis of this overlapping gene set suggests modulation of the nuclear factor of activating T cells 5 (*NFAT5*), *SP1*, *IL4*, *Tnf,* and VEGF pathways, which results in downstream enhancement of innate immune cell responses, Figure 7B. Furthermore, direct immunoblot analysis of protein expression in lung tissue homogenates from wt mice demonstrates enhanced expression of RECK and decreased expression of MMP14 in rTIMP2-treated and tumor-free compared with vehicle control treated (HBSS) mice. In addition, these results demonstrate a trend towards decreased expression of Fibronectin 1, *Myc*, and *Twist* in the tumor-free and rTIMP2-treated wt mice.

In our final experiment, we examined the effect of rTIMP2 treatment on the metastatic niche microenvironment utilizing a syngeneic, heterotopic model with subcutaneous implantation of LL2-luc2 cells in the dorsal flank of wt C57Bl/6 mice, Figure 7C. The results of these experiments again demonstrate reduced growth of primary (heterotopic subcutaneous) tumors in mice treated with rTIMP2 starting on either Day 1 or 7. Although RNAseq analysis of the lung tissues from these mice demonstrated no significant difference in the metastatic tumor burden between control and rTIMP2-treated mice, the expression of *Klf4*, a premetastatic niche marker (*36*), was significantly reduced in both cohorts of rTIMP2 mice. Further gene set analysis of both treatment cohorts suggests there is a downregulation of multiple cell stress-related genes, including multiple members of the heat shock protein (HSP) family, consistent with an overall suppression of the heat shock protein response.

Here we directly demonstrate the tumor suppressor and potential antimetastatic activity of TIMP2 in murine models of NSCLC. In addition, we empirically identified that the loss of function mutation of TIMP2, which eliminates the α3β integrin binding domain, could prime myeloid precursors of the inflammatory response in the absence of pathologic insult, as suggested by the upregulation of *Myadm* expression in non-tumor bearing mT2 mice. However, during the pathologic changes associated with tumor growth, this prior myeloid priming by *Myadm*, observed in non-tumor-bearing mT2 mice, could initiate the enhanced MDSC response observed in tumor-bearing mT2 mice. This premise is consistent with a previous report that demonstrates that *Myadm* expression is positively correlated with the infiltration of MDSCs and is associated with a poor prognosis in NSCLC (*29*). This premise is in line with our proposed role of TIMP2 as a negative regulator of MDSCs (*31*).

Curiously, the functional mutation of the *Timp2* gene did not induce a significant upregulation of other *Timp* genes, suggesting that these other family members did not compensate for the *Timp2* loss of function. The diminished scope of gene changes associated with the *Timp2* mutation is consistent with the lack of phenotypic changes in these mice. However, significant transcriptome changes were observed in both the mT2 mice and their wt littermates <u>following</u> tumor inoculation of mice with the LL2-luc2 tumor cells. Inspection of these transcriptomic changes suggests that the responses in mT2 and wt mice are divergent. However, a closer study reveals that the changes following rTIMP2 treatment are more restricted in mT2 mice (236 genes) compared with those in wt mice (2020 genes). Interestingly, Ingenuity Pathways Analysis of the shared gene changes between rTIMP2-treated wt and mT2 orthotopic tumor models (119 genes) suggests that rTIMP2 treatment functions in both mT2 and wt mice to modulate MDSC responses while enhancing T cell functions. These rTIMP2-mediated gene changes may facilitate responses similar to those mediated by immune checkpoint inhibitors, such as PD-1 and PD-L1 targeting therapeutics that result in reduced inflammatory cell suppressor activity and enhance innate immune responses. In contrast, our studies of the response to rTIMP2 treatment in wt mice suggest that there is also a disruption of oncogenic drivers that can directly influence tumor cell responses. This implies that in wt mice, without the *Myadm* primed myeloid precursors observed in mT2 mice, the effect of rTIMP2 treatment targets both the immune cell responses as well as directly altering tumor cell proliferative (growth) responses.

To conclude, we present new evidence of the anti-tumor capabilities of TIMP2. Mice harboring a loss-of-function mutation in *Timp2* demonstrate a marked increase in tumor growth, and this effect can be rescued with exogenous rTIMP2. Daily intraperitoneal injection of rTIMP2 produces observable benefits in both wt and mT2 tumor models. The success of current treatments for solid malignancies is heavily influenced by the tumor microenvironment (TME). This is exemplified by the promising outcomes following immunotherapy in many solid tumors. We propose that rTIMP2, or functional derivatives thereof, acting as non-cytotoxic agents, may support the efficacy of current and next-generation solid cancer therapies through normalizing the gene expression pattern towards that observed in normal, non-cancerous microenvironment. The results of the present study suggest that rTIMP2 treatment facilitates an anti-tumor host response that involves cellular elements of the tumor microenvironment, namely both immune and endothelial cell functions, as well as direct suppression of tumor cell oncogenic signaling.

This normalized microenvironment is also reflected in the downregulation of the cell stress response (Hsf-regulated genes), reduced tumor vascularity and permeability, potentially enhancing drug delivery and improving treatment responses in patients with NSCLC. Our findings support the emerging concept that the restoration of both the composition and structural integrity of the microenvironment to a pre-malignant state (“normalization of the ECM) can facilitate and enhance cancer therapies.

## MATERIALS AND METHODS

### Study design

The principal objective of this study was the characterize the effects of TIMP2 directly on tumor growth and metastatic niche formation in a murine model of NSCLC and analysis of associated changes in gene expression. We performed *in vivo* studies using both wt and mT2 mice, initially with ten mice of each genotype were randomized to each treatment group as this group size yielded statistically significant data for a similarly designed study in a model of triple-negative breast cancer (TNBC) (*20*). All *in vitro* studies were conducted using samples from 5 mice per group (immunoblot analysis) or assayed in triplicate. Data analysis was performed in an unblinded manner.

### Orthotropic mouse model of lung cancer

For intratracheal instillation, 1× 10^6^ LL2-luc2 cells in 50µL PBS/mouse were intratracheally administered as previously described (*49*). Briefly, mice were anesthetized by intraperitoneal injection (IP) injection with 0.1mL of a mixture of 10mg/mL ketamine and 1mg/mL xylazine diluted in sterile PBS and then placed on an intubation platform. The mouth was opened with tweezers and the trachea was identified using the vocal cords for guidance while holding the tongue. The tumor cell-containing PBS was installed into a predetermined site, at the opening of the trachea without damaging the surrounding tissues. Mice were monitored for tumor progression by bioluminescence in vivo imaging.

### Subcutaneous tumor metastasis model

1× 10^6^ LL2-Luc cells in 100µL PBS/mouse were subcutaneously injected at the dorsal flank, and tumor progression was assessed by caliper measurements and/or *in vivo* bioluminescent imaging.

### In vivo tumor imaging

Tumor growth was monitored by bioluminescence imaging using the Xenogen IVIS Spectrum Imaging System (Perkin Elmer). For luminescence measurements, mice were intraperitoneally injected with 100uL 30mg/mL Luciferin solution (D-Luciferin Firefly Potassium, Caliper Life Sciences) 15 minutes before imaging. The images were obtained with times ranging from 1-120 seconds, depending on the levels of luciferase activity. Living Image Software v3.1 was used for analysis (Caliper Life Sciences).

### Antibodies: See Table S3

Tumor microvascular density was assessed using the rat anti-mouse CD31 (PECAM-1, clone SZ31) antibody (Dianova). The following antibodies were used to characterize immune cell subpopulations in the LL/2 tumors: rat anti-CD45 (Novus Biologicals), rabbit anti-CD11 b, rat anti-Ly6G/Ly6c (Novusbio), mouse anti-CD3, rat anti-CD4, goat anti-CD8 (Santa Cruz).

### RNA and protein purification

Lung tissues from healthy and tumor-bearing mice were harvested and immersed in RNAlater solution (Invitrogen). Alternatively, snap-frozen tissues and OCT-embedded tissues were thawed for 24 hours at -20°C in RNAlater-ICE solution (Invitrogen). RNA and protein were harvested from the same sample using a modification of the Nucleospin RNA/Protein kit (Machery-Nagel). In this modified protocol we included 0.5% antifoam solution in the sample homogenization which was performed using Y-30 using M-tubes (Miltenyi Biotech), and a GentleMACS disruptor on the “RNA” protocol (Miltenyi Biotech). These lysates were passed through a 20-ga needle at least 5 times to produce the stock lysate. Stock lysates were then processed for total RNA and protein isolation following the manufacturer’s instructions (Machery-Nagel). RNA samples were quantified and quality-checked by 280 nm absorbance and TapeStation gel electrophoresis (Agilent) before use. For total protein isolation, a final centrifugation step at 11000 x G for 1 minute was performed before the supernatant was collected, quantified using the Protein Quantification Assay (Machery Nagel), and aliquoted.

### Flow cytometry

After harvesting the lungs were minced with scalpel blades and incubated for 30 min at 37 °C in Dispase (Corning Inc.). The lung tissue fragments were next incubated for 45 min at 37 °C in RPMI 1640 (Invitrogen) containing 0.01% DNAse I (Sigma-Aldrich) and, respectively, 0.25 or 0.17 mg ml^−1^ Liberase (Roche Diagnostics). The digested lungs were passed through a 1 ml syringe to make single-cell suspensions. The cells were filtered through 70 μm nylon mesh and washed before staining. The antibodies used for flow cytometry with mouse cells are listed in Supplementary Table 1 and were used according to the suppliers’ specifications. AccuCount Fluorescent particles (Spherotech) were used for counting the cells.

The analysis of cells after staining was done using an LSR-II flow cytometer (BD Biosciences), and the data were analyzed using FlowJo software (BD Biosciences). Each flow cytometry plot shows cells prepared from a single lung, and aggregate data for each genotype or treatment were used for statistical analysis (median ± 95 % confidence interval (CI) and summary data plots.

### RNA sequencing

RNA quality was assessed using a TapeStation system (Agilent), and all samples received a RIN score >6 before bulk RNA sequencing analysis. PolyA selection was used to enrich for mRNA and RNA sequencing was performed by Azenta Life Science. FASTQ files were analyzed using Partek Flow Software. Read contaminants and low-quality reads (Phred score <20) were removed using Bowtie 2. Trimmed reads were aligned to the mouse (mm10) genome using STAR and annotated to RefSeq transcripts using Partek’s expectation-maximization algorithm. Counts were subjected to median ratio normalization before cluster analysis and differential expression analysis (DEseq2).

### Statistical analysis

All statistical analyses were performed using GraphPad Prism version 10. For comparison of two groups, two-tailed unpaired *t*-tests were used. Kaplan-Meier survival data were analyzed using a log-rank (Mantel-Cox) test. Data are expressed as mean ± SEM, or median ± range (95% CI), of *n* experimental data points. *P* values less than 0.05 were considered significant.

## Acknowledgments

The authors thank the NCI Histopathology Laboratory in Frederick, MD for assistance with histochemical staining and Dr. John J Caterina, NIDCR, NIH for supplying the mutant *Timp2* mice on the original genetic background (C57BL/6j and 129/ReJ) (*21*). Graphics and illustrations for figures were generated using BioRender.com (Toronto, ON, Canada).

## Funding

This work is supported by the Center for Cancer Research (NCI Intramural Program), National Cancer Institute, National Institutes of Health grants ZIA BC011204 & ZIA SC 009179 (WGSS).

## Authors’ Contributions

Conceptualization: SK, DP, SMJ, WGSS

Methodology: SK, DP, TPS, YL, AC, SMJ, BW, WGSS

Investigation: SK, DP, TPS, YL, SJM, SCP, JR, SG, WGSS

Writing-original draft: SK, DP, SMJ, WGSS

Writing-review & editing: DP, SK, TPS, SMJ, WGSS

## Data and materials availability

All data associated with this study are present in the paper or the Supplementary Materials. RNAseq data have been deposited in the GEO database (awaiting assignment confirmation).

## Competing Interest

The authors have no potential conflicts of interest to disclose.

## Supplementary Materials

## Supplementary Methods

### Derivation of congenic Timp2 mutant mice (mT2)

T*imp2* mutant mice were first developed in a mixed genetic background (C57BL/6j and 129/ReJ) (*21*). We created a new congenic strain of mutant *Timp2* mice (mT2) containing a targeted deletion of exons 2 and 3 of the *Timp-2* locus in a clean C57BL/6j genetic background. Marker-assisted breeding was utilized to generate the new strain by backcrossing original male mT2 mice (C57BL/6j and 129/ReJ) with wt C57BL/6j females purchased from The Jackson Laboratory. Ten male carriers of the *Timp2* mutation were selected after each backcross and subjected to full genome scans using 96 polymorphic microsatellite markers spanning all autosomal chromosomes. Mice with the highest number of homozygous C57BL/6j markers were selected as breeders for the next generation. mT2 mice were considered congenic when all 96 microsatellite markers were homozygous to C57BL/6j. The deletion of exons 2 and 3 was confirmed by PCR, as reported previously (*21*). Sequences for the wt-specific and mT2-specific primer pairs are identical to those previously reported (*21*). Mice were housed 5 mice per cage, under a 12-hour light/dark cycle with access to Purina chow and water *ad libitum.* All animal procedures reported in this study were performed by CCR staff. All staff and protocols were approved by the NCI Animal Care and Use Committee (ACUC, ASP # LP-003-4) and followed federal regulatory requirements and standards. All components of the intramural NIH ACU program are accredited by AAALAC International procedures were approved.

#### Cell culture

Lewis lung carcinoma cells stably expressing the Luc2 Luciferase gene (LL/2-Luc2) were obtained from the American Type Culture Collection (ATCC) and cultured in Dulbecco’s Modified Eagle Medium (DMEM), 10% fetal bovine serum (FBS), 1% GlutaMAX, 1% penicillin-streptomycin (P-S). LL/2 spheroids were generated at 5000 cells per well using ultra-low attachment round-bottomed 96-well plates (Corning) and cultured for 6 days to allow efficient spheroid formation. Primary murine lung fibroblasts were cultured in Eagle’s Minimum Essential Medium, 10% FBS, 1X non-essential amino acids (ThermoFisher), and 1% P-S. An automated cells counter, LUNA-FL (Logos Biosystems Inc.), was used to determine the viability, and cell cultures with greater than 95% viability were used for experiments.

#### Preparation of recombinant TIMP2

rTIMP2 with a C-terminal 6x-Histidine tag was produced as previously described (*50*). Lyophilized rTIMP2 was resuspended in sterile Hank’s balanced salt solution (HBSS) at a concentration of 6ug/100uL and stored at -80° C in single-use aliquots ready for intraperitoneal injection.

#### Lung fibroblast isolation

Murine lung tissues were minced using two scalpels, then transferred to a 30mL Erlenmeyer flask with 10mL DMEM-F12 media supplemented with 10mM HEPES, 200U/mL collagenase D, 2.4U/mL Dispase, 100U/mL DNA I, and 1% penicillin-streptomycin (P-S). The digestion mix was incubated at 37°C, stirring slowly, for approximately 40 minutes (when lung fragments changed color from red to white, forming sticky fibers). 40mL DMEM-F12 10% FBS 1% P-S (DMEM-F12 FM) was added to the digestion mix and mixed extensively by pipetting to resuspend the fragments. The tissue fragments were washed 3x with 20mL DMEM-F12 FM, pelleting the tissue at 500G for 5 minutes in between washes. Tissue fragments were resuspended in 10mL DMEM-F12 FM and plated in 10cm dishes. Stromal cells (fibroblasts) exit the tissue fragments over the next few days, attach to the tissue culture plastic, and begin to proliferate. Between days 7-14, the tissue fragments were washed away, and culture media was changed to EMEM 10% FBS, 1X non-essential amino acids, and 1% P-S to support fibroblast propagation.

#### Immunoblot analysis

Tissue homogenates containing TCEP were boiled for 3 minutes and loaded into precast 4-20% polyacrylamide gels (Bio-Rad). SDS-PAGE was performed, and proteins were transferred to nitrocellulose membranes using a Trans-Blot Turbo system (Bio-Rad). The membranes were blocked for 1 hour in 2.5% milk TBS 0.1% Tween 20, before immunostaining. The antibodies used are summarized in Supplementary Table 1. Either alpha-tubulin or total protein (Bio-Rad stain-free gels or Ponceau stain) was used to normalize sample loading.

### Immunohistochemistry & Immunofluorescence

Immunohistochemistry (IHC) and immunofluorescence (IF) were performed on formalin-fixed paraffin-embedded (FFPE) or frozen tissue embedded in optimal-cutting temperature (OCT) medium using standard laboratory practices for processing, embedding, sectioning, and staining. Primary antibodies are listed in Table S3.

### Supplementary Material

**Figure S1.**
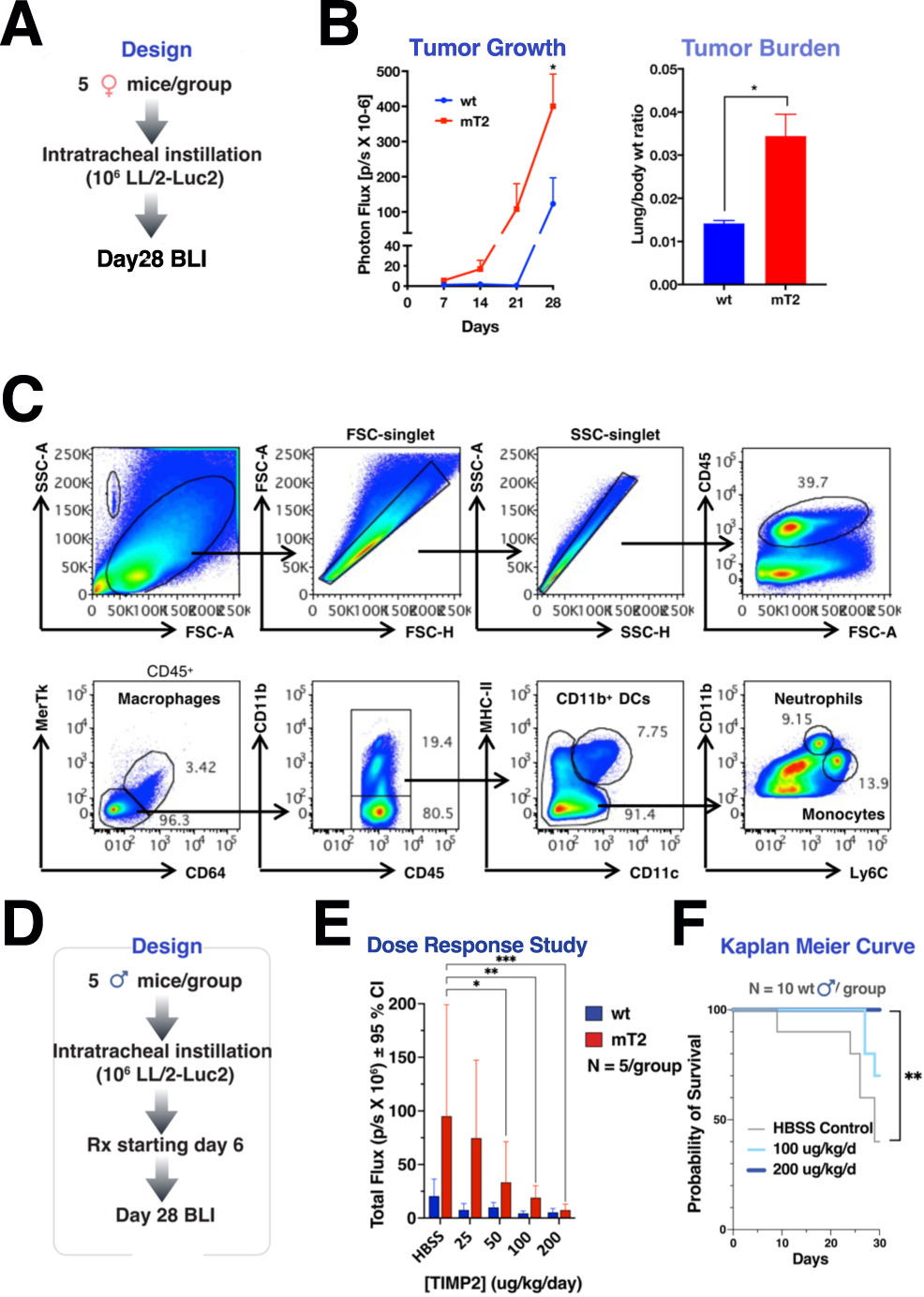
(A) Experimental design in a female-only study comparing wt vs mT2 orthotopic tumor growth. (B) Metrics describing the differences in orthotopic LL/2-Luc2 tumor growth between wt and mT2 tumor-bearing female mice. (C) The gating strategy utilized to compare myeloid cell populations in wt and mT2 mice. (D) Design of a pilot orthotopic tumor study investigating the effective dose range for rTIMP2, and (E) in vivo imaging depicting differences in tumor volume across the treatment regimens. (F) Kaplan-Meier survival analysis comparing untreated, 100ug/kg/day, and 200ug/kg/day rTIMP2 treatment regimens in wt mice.

**Figure S2.**
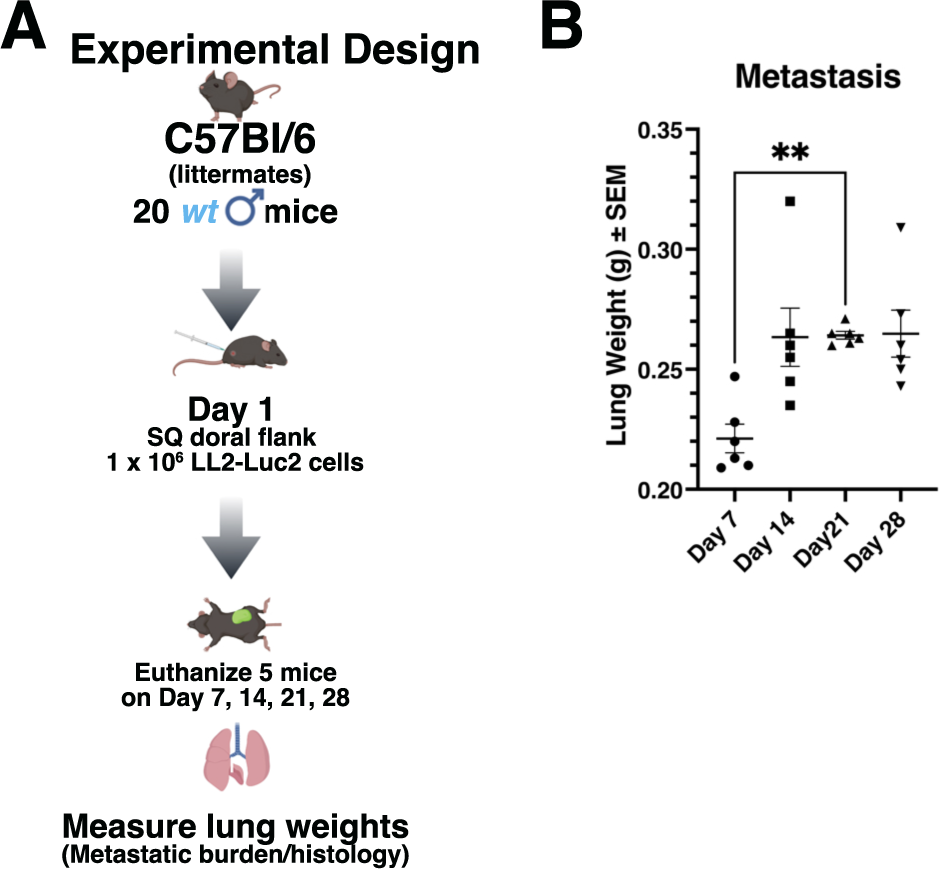
(A) Experimental design in study to examine time frame of lung metastasis from dorsal flank tumors in wt mice. (B) Lung weights (correlating with metastatic burden) of wt mice during the time frame to metastasis outlined in panel A. Data shows that the period between day 7 and day 21 post-tumor inoculation is critical for metastasis formation and development of the metastatic niche.

**Table S1. Summary of orthotopic Lewis Lung Carcinoma RNA sequencing data and gene set analysis using Ingenuity Pathway Analysis.**

**Table S2. Summary of subcutaneous heterotopic Lewis Lung Carcinoma RNA sequencing data and gene set analysis using Ingenuity Pathway Analysis.**

**Table S3. Antibodies and reagent list.**

**ARRIVE Author Checklist**

## Notes

### Competing Interest Statement

The authors have declared no competing interest.

